## Supplemental Figures for "Down-regulation of AKT proteins slows the growth of mutant-KRAS pancreatic tumors"

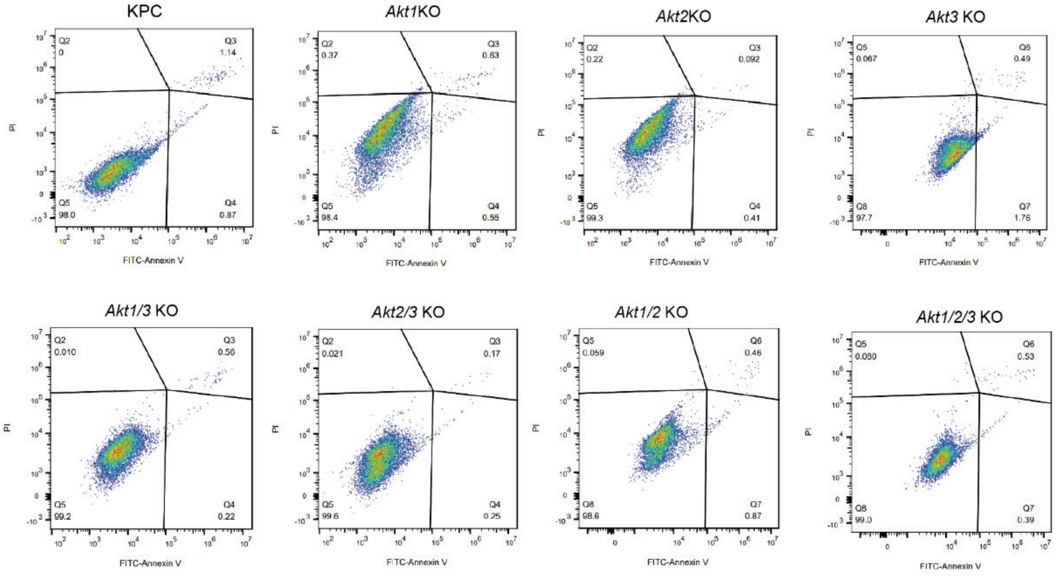


**Supplemental Figure S1.** KPC and *Akt1/2/3*KO cells show comparable apoptotic rates. Apoptosis assay by ANNEXIN-V and PI staining and flow cytometry analysis. The cells were set up as the cell counting assay in Figure 3(A) and evaluated on day 4 by flow cytometry, representative images of technical replicates (n=2) are shown.

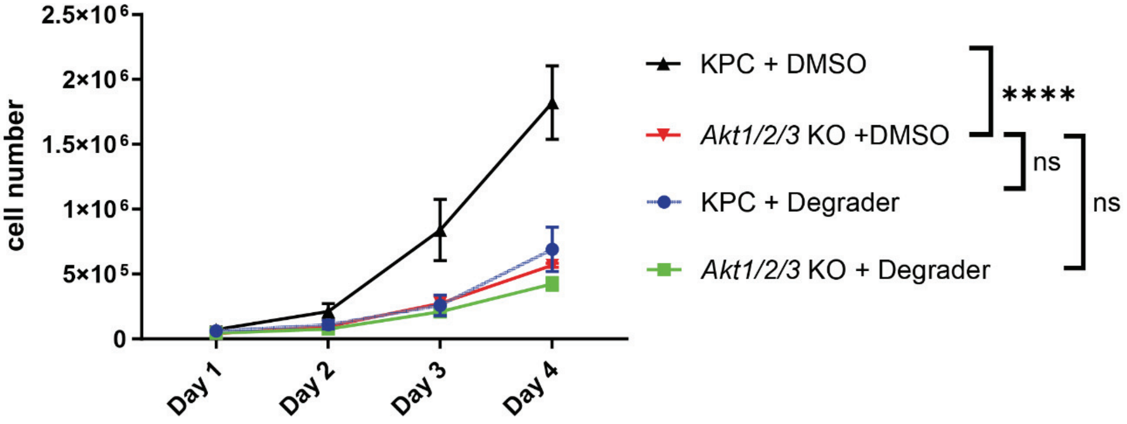


**Supplemental Figure S2**. AKT Degrader phenocopies *Akt1/2/3* genetic ablation in impeding cell growth. 50,000 cells were seeded in triplicates in each well of 6-well plates in 5% FBS DMEM with DMSO, or 500nM AKT Degrader. Cells were counted over the following 4 days. The experiment was repeated three times. Two-way ANOVA was performed, followed by uncorrected Fisher's LSD multiple comparisons of the indicated pairs. ****$p<0.0001$

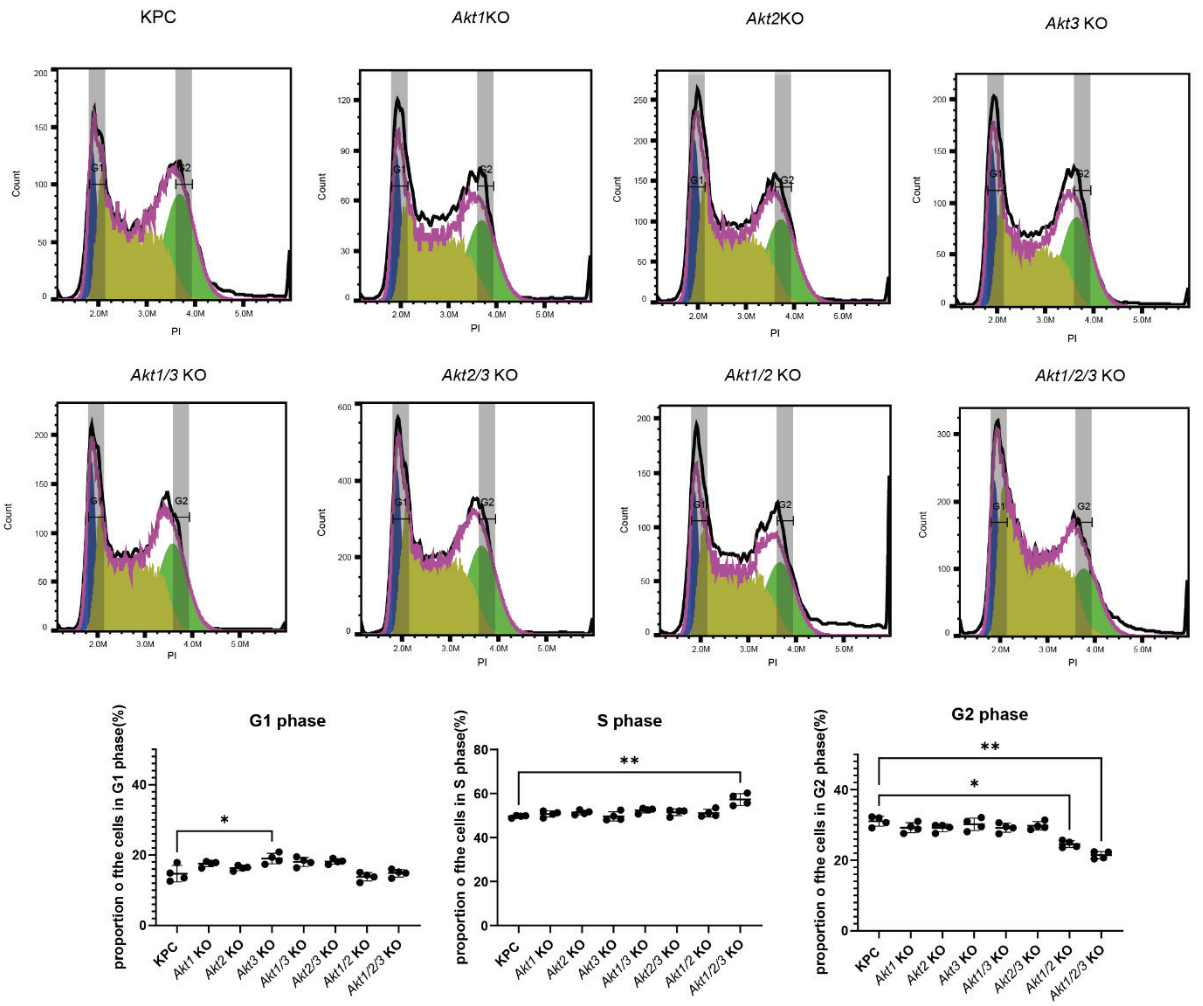


**Supplemental Figure S3.** KPC and *Akt1/2/3*KO cells show comparable cell cycle distribution. Cell cycle analysis based on PI staining of the nuclear DNA. The cells were set up as in Figure 3(A) in technical replicates (n=4), and evaluated on day 2 by flow cytometry. Kruskal–Wallis ANOVA (nonparametric) tests were performed comparing proportions of G1 (P=0.0019), S (P=0.0112) and G2 (P=0.0051). Dunn's multiple comparisons test was made between KPC and each of the other lines, but only statistically significant comparisons are shown. * P < 0.05, ** P < 0.01, *** P < 0.001, **** P<0.0001.

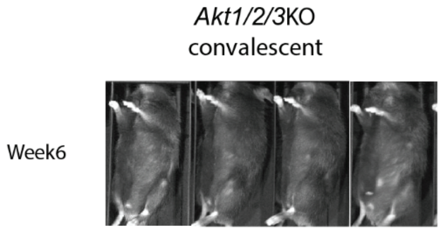


**Supplemental Figure S4.** *Akt1/2/3*KO cells-implanted mice have long-term survivors. Representative IVIS images of all four tumor-free long-term survivors implanted with *Akt1/2/3*KO cells at week 6 post-implantation.

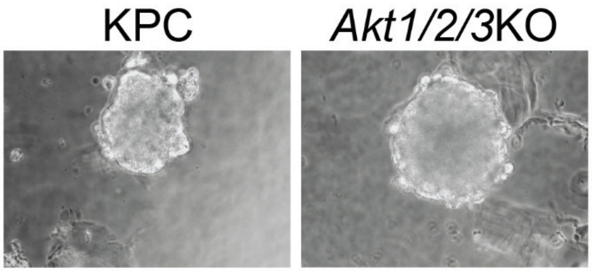


**Supplemental Figure S5.** Both KPC and *Akt1/2/3*KO cells are capable of anchorage-independent growth. 3D culture in 0.24% methylcellulose 10%FBS DMEM indicates that both KPC and *Akt1/2/3* KO cells are capable of anchorage-independent growth. Pictures were taken at 40X magnification with an inverse microscope.

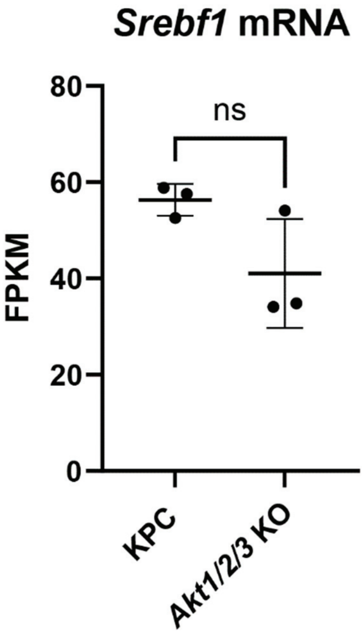


**Supplemental Figure S6.** The transcript level of *Srebf1* in KPC versus *Akt1/2/3*KO cells.

A

B

C

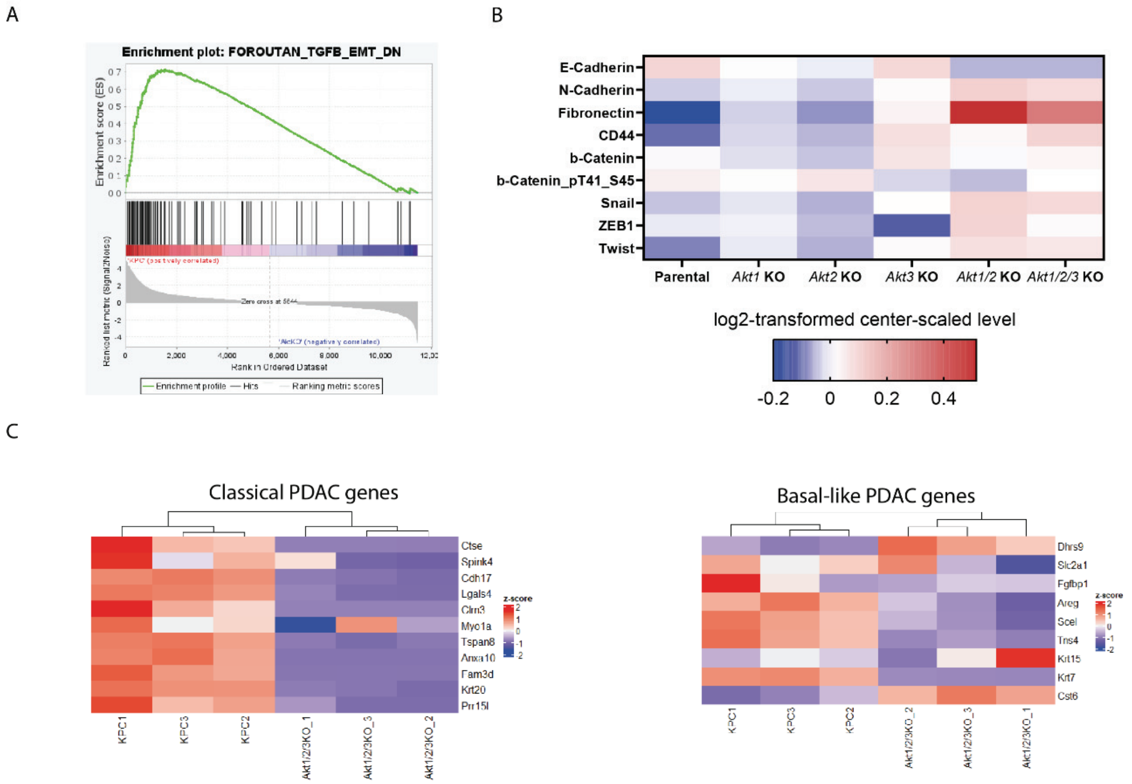


**Supplemental Figure S7.** RNA-seq and RPPA suggest *Akt/1/2/3*KO cells undergo epithelial-to-mesenchymal transition (EMT). (A) Representative EMT gene set in GSEA pathway enrichment analysis. (B) Relative protein levels by RPPA of EMT-signature genes of the indicated groups. (C) Heat maps comparing KPC and *Akt1/2/3*KO cells in terms of human PDAC subtypes signature genes.
